# “A human *CHRNA6* 3′ UTR variant alters the molecular environment and microRNA engagement of the adolescent VTA without changing *Chrna6* transcript abundance”

**DOI:** 10.64898/2026.10.09.758057

**Authors:** Yasamin Hajy Heydary, Emily M. Castro, Kaylie S. Gurholt, Nikki Le, Frances M. Leslie, Shahrdad Lotfipour

## Abstract

A single nucleotide polymorphism (rs2304297) in the 3′ untranslated region (UTR) of *CHRNA6*, encoding the α6 nicotinic acetylcholine receptor subunit, is associated with adolescent smoking behavior. Humanized rats carrying the “C” or “G” allele show sex- and genotype-dependent responses to adolescent nicotine, yet the variant does not alter the protein sequence and the mechanism has remained unknown. We examined the molecular consequences of rs2304297 *in vivo* in the adolescent ventral tegmental area (VTA). *Chrna6* transcript abundance did not differ by genotype, nor did transcript stability or polyadenylation site selection. The variant was instead associated with a transcriptional signature spanning myelin maintenance (*Enpp6*), lipid metabolism (*Retsat*), and genomic repair (*Polq*), and with elevated basal lipid peroxidation in α6^GG^ animals. Computational folding predicted greater conformational flexibility of the “G” allele, and spatial colocalization of *Chrna6* with miRNA-365-5p was reduced in α6^GG^ animals of both sexes. Coordinated expression of the two transcripts, however, followed a sex-by-genotype pattern, absent in α6^CC^ females and α6^GG^ males. These same groups showed elevated tyrosine hydroxylase expression and dopaminergic neuron number, and expansion of the *Chrna6* co-expression network following nicotine. We propose that rs2304297 acts through two distinct layers: genotype determines spatial engagement of miRNA-365-5p with the transcript, while sex determines whether that engagement produces functional regulation. These findings identify a post-transcriptional mechanism for a non-coding nicotine addiction risk variant and demonstrate that its molecular effect can be uniform while its consequences are not.

## Introduction

Adolescence is a period of extensive neurodevelopmental reorganization during which reward and decision-making circuits undergo synaptic refinement, while prefrontal inhibitory control matures relatively late [1–2]. Nicotine exposure within this window disrupts maturing neurotransmitter systems, produces lasting alterations in reward circuitry, and elevates the risk of subsequent substance use, including illicit stimulants such as cocaine and methamphetamine [3–6]. Tobacco use among the youth remains a pressing concern with more than 2.25 million U.S. middle and high school students reported current tobacco product use in 2024, driven largely by the accessibility of electronic cigarettes [7]. Despite this, the molecular mechanisms that render the adolescent brain selectively susceptible to nicotine remain poorly defined.

Nicotine exerts its reinforcing effects through neuronal nicotinic acetylcholine receptors (nAChRs), pentameric ligand-gated ion channels assembled from α and β subunits [8–9]. Among these subtypes, α6-containing nAChRs (α6*nAChRs) have a distinctive position in addiction pathology. Their expression is largely restricted to catecholaminergic nuclei, including the mesolimbic dopamine pathway projecting from the ventral tegmental area (VTA) to the nucleus accumbens (NAc), they reach peak expression during adolescence. Pre-clinical work has established that they are required for nicotine-induced locomotor activity, dopamine release, and self-administration [10–13]. Individual susceptibility to nicotine dependence is substantially heritable [14–15]. Clinical studies have repeatedly linked rs2304297, a single nucleotide polymorphism in the 3′ untranslated region (UTR) of *CHRNA6*, to adolescent smoking behavior, with the G allele treated as the risk variant [16–19].

To test the functional consequences of this variant *in vivo*, our laboratory generated a humanized rat model in which the endogenous 95 bp rat *Chrna6* 3′ UTR was replaced with the corresponding 559 bp human sequence carrying either the C or G allele, producing lines homozygous for each (α6^CC^ and α6^GG^) [20]. These animals show no baseline behavioral differences, yet sub-chronic nicotine exposure produces sex- and genotype-dependent divergence. Of note, there is increased locomotor and anxiolytic behavior in α6^CC^ females and α6^GG^ males, accompanied by corresponding shifts in nucleus accumbens shell dopamine overflow and metabolite profiles [20–21]. Because rs2304297 is located outside the coding sequence, it cannot alter the amino acid sequence of the α6 subunit. Any functional consequence is likely exerted through the regulatory capacity of the 3′ UTR, which governs transcript stability, subcellular localization, translational efficiency, and susceptibility to microRNA-mediated repression [22].

Several 3′ UTR variants are known to influence human disease risk through exactly these mechanisms. The rs10954213 variant in the *IRF5* 3′ UTR strengthens a proximal polyadenylation signal, shortening the transcript and increasing protein output in a manner that confers autoimmune risk [23]. The rs12720208 variant in the *FGF20* 3′ UTR disrupts a miR-433 binding site, releasing translational repression and driving downstream α-synuclein pathology in Parkinson’s disease [24]. Within the mesolimbic system specifically, the variable number tandem repeat in the *SLC6A3* 3′ UTR alters dopamine transporter density and constitutes an established risk factor for substance use disorders [25]. None of these possibilities has been tested for rs2304297 *in vivo*.

We therefore asked how a non-coding variant that leaves the α6 protein sequence intact produces measurable behavioral and neurochemical consequences, and why those consequences depend on sex. Using bulk RNA sequencing of the adolescent VTA, we first mapped the transcriptomic landscape associated with genotype, sex, and sub-chronic nicotine exposure, and tested the physiological relevance of the resulting metabolic signature by quantifying tissue lipid peroxidation. We then evaluated the post-transcriptional routes available to a 3′ UTR variant: allele-specific secondary structure, transcript stability, alternative polyadenylation site selection, and the organization of the *Chrna6* co-expression network. Finally, we examined microRNA-365-5p, a conserved candidate predicted to bind the *CHRNA6* 3′ UTR adjacent to the polymorphic site, at both the expression and spatial levels, and assessed the resulting dopaminergic phenotype by quantifying tyrosine hydroxylase expression and dopaminergic neuron number in the VTA. Collectively, these experiments reveal the regulatory level through which rs2304297 acts and clarify the emergence of its sex-dependent effects.

## Materials and methods

### Animals and Generation of Humanized Rats

All procedures were approved by the Institutional Animal Care and Use Committee (IACUC) at the University of California, Irvine, and conducted in accordance with AAALAC guidelines. Two rat lines with a humanized *Chrna6* 3’ untranslated region (UTR) were generated via CRISPR/Cas9 editing (Cyagen Biosciences), replacing the endogenous 95 bp rat sequence with the 559 bp human sequence containing either the ’C’ or ’G’ allele of the rs2304297 SNP, as previously detailed (Cardenas et al., 2022). The *CHRNA*6 3′-UTR SNP rats (SD-*CHRNA*6e^m1Slot^ ^984^, SD-*CHRNA*6^em2Slot^ ^985^) have been shared and stored at the Rat Resource and Research Center (RRRC). Rats were weaned at postnatal day 21 (P21), group-housed by sex under a 12-hour light-dark cycle, and provided *ad libitum* access to food and water. Experimental groups were balanced using no more than one male and one female per litter to control for litter effects. For drug-naive experiments, tissue samples were collected on P31.

### Surgical Procedures and Sub-Chronic Nicotine Administration

At P24, adolescent rats were anesthetized with Equithesin (0.035 mg/kg, i.p.) and implanted with a right jugular vein catheter [26]. Carprofen (4 mg/kg, s.c.) was administered for postoperative analgesia. Following a three-day recovery period (P25-P27), rats received sub-chronic nicotine treatment from P28 to P31. Animals received daily pretreatments of two intravenous (i.v.) injections of either nicotine (30 µg/kg per injection, calculated as free base, pH 7.2-7.4) or sterile saline, administered one minute apart. On P31, thirty minutes post-injection (representing the half-life of nicotine), animals were rapidly decapitated. Brains were extracted, flash-frozen in 2-methylbutane (−20°C), and stored at -80°C until further processing.

### Tissue Processing and RNA Extraction

Brain tissues were sectioned at 300 µm on a cryostat. VTA tissue punches (1-mm) were collected using standardized anatomical landmarks. For RNA sequencing (RNA-seq) and associated validations, total RNA was extracted using the PureLink RNA Mini Kit (Invitrogen). For microRNA (miRNA) analyses, RNA was extracted using the miRNeasy Tissue/Cells Advanced Micro Kit (Qiagen). RNA concentration and purity were evaluated via NanoDrop 2000c (Thermo Fisher Scientific). RNA Integrity Number (RIN) was assessed using an Agilent 2100 Bioanalyzer (UCI Genomics Research & Technology Hub) and samples with RIN < 7.5 were excluded.

### RNA Sequencing and Bioinformatics Analysis

Illumina sequencing libraries were prepared by the UCI GRT Hub and sequenced across three batches, yielding 150 bp paired-end reads (n=9 per group; total N=72). Raw read quality was assessed with FastQC (v0.11.9) and trimmed using Trimmomatic (v0.39) [27]. Trimmed reads were aligned to a custom rat reference genome, incorporating the humanized *Chrna6* locus, using STAR (v2.7.10b) [28]. Alignment fidelity at the *Chrna6* locus was verified using the Integrative Genomics Viewer (IGV). Read quantification was performed with featureCounts (v2.0.3) [29]. Differential expression analysis was executed in R (v4.3.x) using DESeq2 (v1.42.0) [30], incorporating sequencing batch as a covariate. Pairwise contrasts evaluated specific experimental conditions using the Wald test, while two-way and three-way interactions were assessed via the Likelihood Ratio Test (LRT). Significance was defined as an FDR-adjusted *p* < 0.05 and |log2FC| ≥ 1.0. Functional enrichment of synaptic processes was determined using clusterProfiler (v4.10.0) against the SynGO database. Protein-protein interaction (PPI) networks for genes highly correlated with *Chrna6* (|r| > 0.90) were constructed using STRING (v12.0), utilizing a high-confidence threshold (0.700) and excluding disconnected nodes.

### Rapid Amplification of cDNA Ends (RACE) and Isoform Characterization

Alternative polyadenylation (APA) isoforms of *Chrna6* were characterized using the 3’ RACE System (Invitrogen) utilizing a specific forward primer (5’-TGTTCAGAAATTTACAGACACC-3’) binding upstream of the first predicted polyadenylation site. Target amplicons (∼600 bp) were gel-purified (QIAquick Gel Extraction Kit, Qiagen) and cloned into the pJET1.2/blunt vector (CloneJET PCR Cloning Kit, Thermo Scientific). Following transformation into *E. coli*, recombinant plasmids from four isolated colonies per group were confirmed via Sanger sequencing (Azenta Life Sciences).

### Quantitative Real-Time PCR (qPCR)

cDNA was synthesized using the High-Capacity cDNA Reverse Transcription Kit for mRNA, and the TaqMan Advanced miRNA cDNA Synthesis Kit for miRNA targets (Thermo Fisher Scientific). qPCR was conducted on a QuantStudio 7 System (Applied Biosystems) using TaqMan Fast Advanced Master Mix and specific TaqMan Assays. To differentiate mature from precursor *Chrna6* transcripts, a pre-designed assay (Rn00589325_m1) was used for mature mRNA (spanning an exon-exon junction), and a custom assay (AR9HPDV) targeted the intron 5/exon 6 junction for the *Chrna6* pre-mRNA. *Gapdh* (Rn01775763_g1) and miRNA-125-5p served as endogenous controls for mRNA and miRNA targets, respectively. Relative expression was calculated using the 2^-ΔΔCt^ method. Transcript stability was determined using a Relative Stability Index [ΔCt (mature mRNA) - ΔCt (pre-mRNA)].

### RNAscope In Situ Hybridization and Spatial Analysis

Animals were transcardially perfused on P31, and brains were cryoprotected in 30% sucrose, flash-frozen, embedded in OCT, and sectioned coronally at 15 µm. Multiplex in situ hybridization was performed using the RNAscope Multiplex Fluorescent Reagent Kit v2 (ACD). Images of the anterior (aVTA; Bregma -5.30 mm) and posterior VTA (pVTA; Bregma -6.04 mm) were acquired using a Zeiss LSM 980 at the Optical Biology Core Facility of the Developmental Biology Center, a shared resource supported by the Cancer Center Support Grant (CA-62203) and NIH-S10OD032327-01 (63X objective). Spatial colocalization was analyzed across five independent fields of view per section using FIJI. Following background subtraction (Rolling Ball, radius = 15 pixels) and Gaussian filtering, transcripts were segmented via local maxima identification. Colocalization was quantified using an object-based center-of-mass distance algorithm (JACoP plugin, v2.1.4) [31].

### Lipid Peroxidation (MDA) Assay

VTA punches were homogenized in a RIPA buffer supplemented with protease and phosphatase inhibitors, and cleared lysates were collected. Total protein concentration was determined via BCA Assay (Thermo Fisher Scientific). Malondialdehyde (MDA) levels were quantified using a Lipid Peroxidation Assay Kit (Abcam). MDA concentration was normalized to total protein (nmol/mg).

### Western blot

P31 male and female *CHRNA6* 3’-UTR SNP rats were rapidly decapitated, and brains were flash frozen in -30℃ 2-methylbutane. Brains were sectioned into 300µm tissue slices (Leica Biosystems) and bilateral 1mm punches were taken from the entire serially sectioned VTA (ten punches total) and NAc (ten punches total). Tissue was prepared and homogenized in neuronal protein extraction reagent (N-PER, Thermo Scientific) and halt protease inhibitor cocktail, EDTA-free (Thermo Scientific) using motorized pellet mixers (VWR International) on ice. Tissue samples were centrifuged at 10,000 x g for 10 minutes then for 30 minutes at 15,000 x g at 4℃. Protein concentration was determined using a bicinchoninic acid (BCA) assay (Thermo Scientific) and prepared with NuPAGE LDS Sample Buffer 4X (Thermo Scientific) and NuPAGE Sample Reducing Agent (Thermo Scientific). Samples were boiled to 70℃ for 10 minutes and immediately placed on ice. Ten micrograms of protein alongside precision plus protein standards were loaded into NuPAGE 4-12% Bis-Tris Protein Gels (Thermo Scientific) in NuPAGE MES Running Buffer (Thermo Scientific) and bands were separated at 120V for 2 hours using XCell SureLock Mini-Cell (Thermo Scientific). Proteins were then transferred to 0.45 µm nitrocellulose membranes (Thermo Scientific) in cold transfer buffer (0.025M Tris base, 0.19M Glycine, 20% (v/v) Methanol) using a Mini Trans-Blot Cell (BioRad) at 120V for 2 hours. Membranes were incubated in blocking buffer (5% Nonfat Dry Milk, TBS-T) for 1 hour at room temperature. Membranes were cut horizontally according to expected band size and then incubated overnight with corresponding antibodies against GAPDH (Thermo, GA1R Dylight 800, 1:1000), Tyrosine Hydroxylase (TH) (Cell Signaling Technology, 58844, 1:1000), Dopamine Receptor D1 (DRD1) (Proteintech, 17934-1-AP, 1:1000), Dopamine Receptor D2 (DRD2) (Sigma-Aldrich, AB5084P, 1:1000), Dopamine Receptor D3 (DRD3) (Abcam, ab155098, 1:1000), and Dopamine Transporter (DAT) (Abcam, ab184451, 1:1000). Membranes were then washed three times for five minutes each in TBS with agitation and incubated in secondary antibody (for GAPDH, goat anti-mouse, Thermo, 62-6520, 1:1000; for TH, goat anti-rabbit, Thermo, 32460, 1:20, for all other antibodies, mouse anti-rabbit, Cell Signaling Technology, 93702, 1:5000). Membranes were washed and Pierce ECL Western Blotting Substrate (Thermo Scientific) was used for chemiluminescent detection and imaged on a Chemi-Doc gel imager (BioRad). Due to the expected proximity of band sizes for VTA DRD2 and DRD3, membranes were imaged for DRD2 first and then incubated in Restore Western Blot Stripping Buffer (Thermo Scientific) for 10 minutes, washed, and stained as described below for DRD3. Images were analyzed using NIH ImageJ software. The net band intensity (band of interest minus background pixels) was normalized to the loading control, GAPDH. Experimenter was blind to sex and genotype groups during band intensity analysis. (See Supporting Information for all representative blot images).

### Immunohistochemistry

P31 humanized rats were perfused transcardially with ice-cold phosphate buffered saline (PBS) followed by 4% paraformaldehyde (PFA). Brains were dissected and postfixed in 4% PFA overnight in 4℃. The next day, brains were cryoprotected in a 30% sucrose solution. Tissue was then rapidly frozen in -30℃ 2-methylbutane and sectioned coronally at 40µm. Tissue was blocked and permeated in 0.2% Triton X-100, 5% normal donkey serum (NDS) in PBS for 1 hour. Tissue was incubated in primary antibody in incubation buffer (Tyrosine Hydroxylase, Thermo, PA1-4679, 1:1000) overnight at 4℃. Omission of the primary antibody was a negative control. Sections were then rinsed and incubated in secondary antibody (donkey anti-sheep Alexa fluor 647, Thermo, A21448, 1:1000) for two hours at room temperature. Sections were rinsed, mounted onto slides, and cover slipped with Fluoromount-G with DAPI (Invitrogen).

### Microscopy and image analysis

Images were captured at 20x magnification (Keyence, Itasca Illinois). At least three intact sections from anterior to posterior VTA were included in analysis. Using NIH ImageJ, the VTA boundary was outlined in each section according to Paxinos & Watson Atlas. Dopamine neurons within the VTA were determined by colocalization of DAPI and TH labeling and manually counted with ImageJ. The experimenter was blind to sex and genotype groups during both imaging and analysis.

### In Silico RNA Secondary Structure Prediction

Secondary structure dynamics of the *Chrna6* 3’ UTR (C and G alleles) were modeled using RNAfold (ViennaRNA Web Services) [32]. Simulations were performed at 37°C using minimum free energy (MFE), partition function, and centroid structure algorithms with default Turner energy parameters.

### Statistical Analysis

Analyses of qPCR, behavioral, and biochemical data were conducted using JMP Pro (v18). For mRNA expression, ΔCt values were analyzed using three-way Analysis of Variance (ANOVA) assessing the main effects of genotype, sex, and treatment. MDA concentrations were evaluated using a Restricted Maximum Likelihood (REML) linear mixed model with experimental assay round as a random effect, followed by Tukey’s HSD post-hoc test for significant interactions. Statistical significance was defined as *p* < 0.05. Data are presented as mean ± SEM unless otherwise specified.

## Results

### Sex and genotype, not nicotine exposure, is the dominant axis of transcriptional variation in the adolescent VTA

To determine how the rs2304297 variant shapes the molecular environment of the adolescent reward circuit, we performed bulk RNA sequencing on the VTA tissue punches from saline- and nicotine-treated male and female α6^CC^ and α6^GG^ rats (n = 9 per group; N = 72). All libraries passed quality control, with a mean sequencing depth of 30.0 million reads and a mean unique mapping rate of 79.5% against a custom reference genome carrying the humanized *Chrna6* locus. Alignment fidelity across the humanized region was verified directly in the Integrative Genomics Viewer. Because libraries were prepared across three sequencing runs, batch was included as a covariate in all differential expression models. Principal component analysis separated samples primarily by sex, which resolved cleanly along PC2 (19% of variance) (**Fig. 1a**). Separation along PC1 (26% of variance) was less pronounced, with α6^CC^ samples distributed toward the left and α6^GG^ samples toward the right within each sex, though the two genotypes somewhat overlapped (**Fig. 1a**). Genotype therefore contributes to global transcriptomic structure alongside sex, though less discretely than the sex-driven separation.

**Figure 1.**
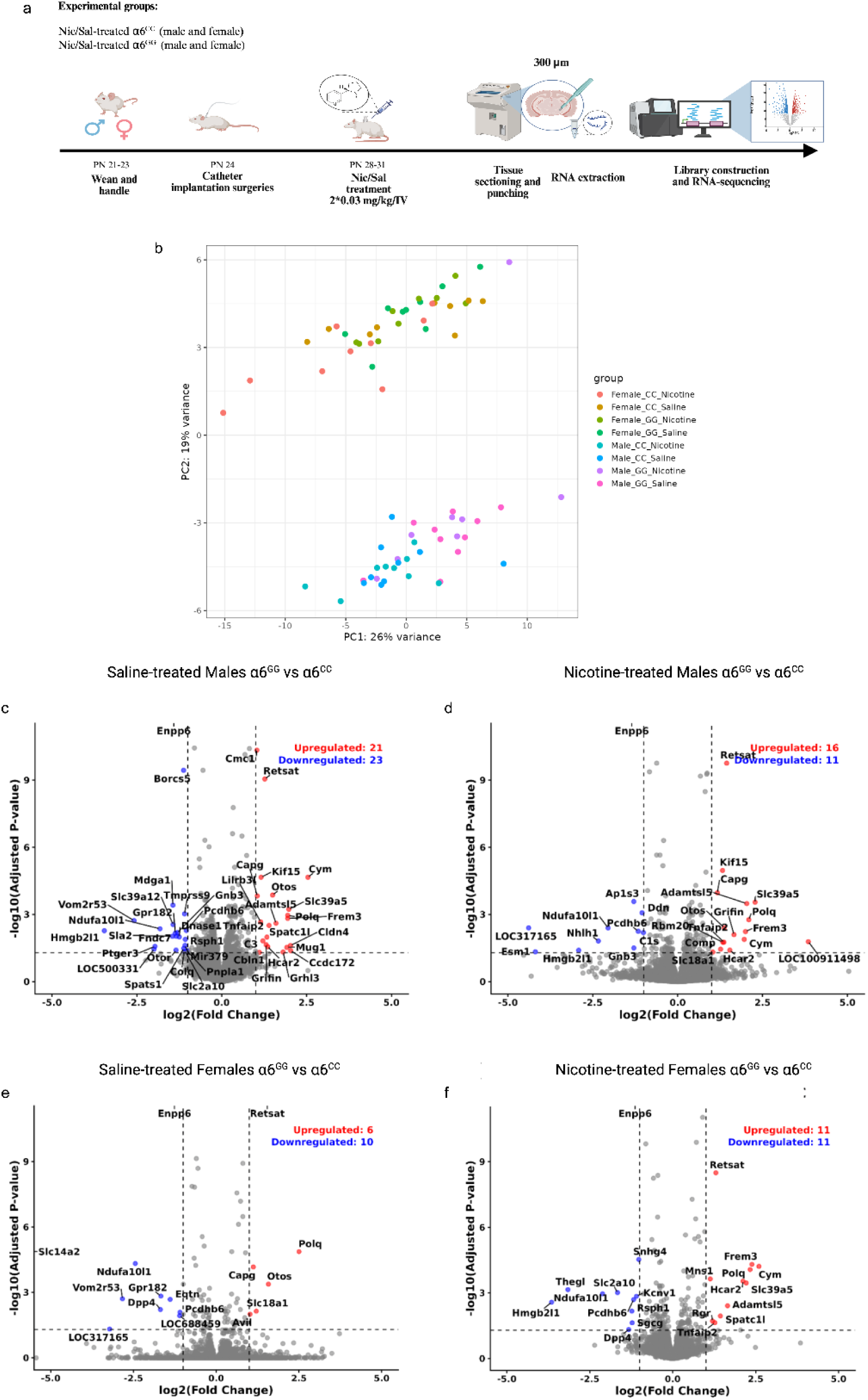
The rs2304297 genotype is associated with a distinct transcriptional profile in the adolescent VTA. **(a)** Experimental timeline. Male and female α6^CC^ and α6^GG^ rats were weaned and handled (PN 21–23), implanted with intravenous catheters (PN 24), and treated with saline or nicotine (2 × 0.03 mg/kg, i.v.) daily from PN 28–31. Brains were sectioned at 300 µm, the VTA isolated by tissue punch, and RNA extracted for library construction and sequencing. **(b)** Principal component analysis of normalized expression across all sequenced samples (n = 9 per group; N = 72), colored by sex, genotype, and treatment. Samples separated by sex along PC2 (19% of variance). PC1 (26% of variance) shows a genotype-associated gradient. **(c–f)** Volcano plots of differential expression between α6^GG^ and α6^CC^ animals within each sex and treatment condition: **(c)** saline-treated males, **(d)** nicotine-treated males, **(e)** saline-treated females, **(f)** nicotine-treated females. Log₂ fold cftange is plotted against −log₁₀ adjusted *P* value. Dashed vertical lines mark log₂ fold change of ±1; the dashed horizontal line marks an adjusted *P* value of 0.05. Red points are upregulated and blue points downregulated in α6^GG^ relative to α6^CC^. Counts of significantly changed transcripts are inset in each panel. Differential expression was assessed by DESeq2 with Benjamini–Hochberg correction. No transcript reached significance for nicotine relative to saline within any sex-and-genotype group.

Direct comparison of nicotine- to saline-treated animals yielded no differentially expressed genes surviving FDR correction in any of the four sex-by-genotype comparisons. At this dose and duration, sub-chronic nicotine exposure does not act as an independent driver of broad transcriptional change in the adolescent VTA. Its molecular signature instead emerges through interaction with genotype. More specifically, when α6^GG^ and α6^CC^ animals are compared separately within saline- and nicotine-treated conditions, different genes and pathways surface as genotype-associated depending on treatment condition (**Fig. 1b–e**). The transcriptional effect of nicotine in this model is expressed through its interaction with genotype rather than independently of it. Moreover, one further observation is central to the rest of this study. *Chrna6* itself was not differentially expressed between genotypes, indicating that the variant does not alter the abundance of the transcript in which it resides. This emphasizes that any functional effect must therefore be exerted through other routes.

### Differential expression identifies a genotype signature in metabolic systems

Stratified comparison of α6^GG^ and α6^CC^ animals identified several genotype-associated transcripts (**Fig. 1b–e**). Three genes were differentially expressed in all four sex-and-treatment comparisons. *Enpp6*, encoding a choline-specific phosphodiesterase required for oligodendrocyte differentiation and myelin maintenance [33], was consistently downregulated in α6^GG^ rats. *Retsat*, encoding retinol saturase and implicated in lipid metabolism and oxidative stress handling [34], and *Polq*, encoding the DNA polymerase that mediates microhomology-mediated end joining of double-strand breaks [35–36], were both consistently upregulated in α6^GG^ rats. Moreover, reduced myelin maintenance alongside increased lipid metabolism and DNA repair points to a tissue under a different metabolic and redox load. Because saline-treated animals of both sexes showed the same pattern, it reflects how the α6^GG^ VTA differs before any nicotine exposure.

We then tested reproducibility in an independent cohort of animals, we included the metabolic and genomic targets identified above and assessing each by three-way ANOVA with sex, genotype, and treatment as factors (**Fig. 2a-c**). *Retsat* showed the strongest genotype effect (F(1,24) = 28.32, p < 0.0001), accompanied by significant genotype main effects for *Enpp6* (F(1,24) = 14.12, p = 0.001), *Polq* (F(1,24) = 9.76, p = 0.005). Independent biological replication of *Retsat*, *Polq*, and *Enpp6* establishes a reproducible rs2304297 signature in the adolescent VTA spanning lipid metabolism, genomic maintenance, and myelination.

**Figure 2.**
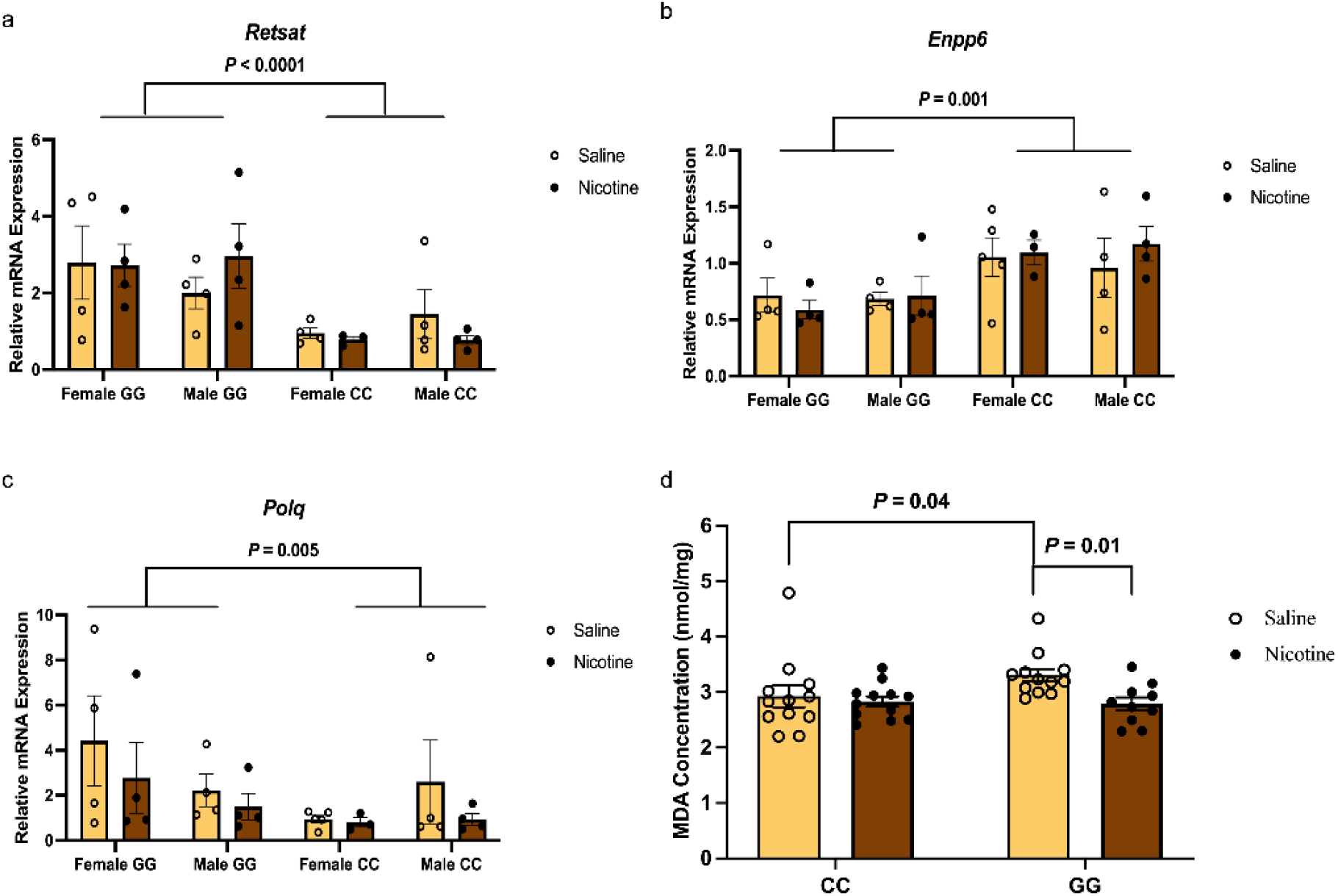
A genotype signature in lipid and genomic maintenance genes is accompanied by elevated lipid peroxidation in α6^GG^ animals. **(a–c)** Relative mRNA expression measured by qPCR in a cohort of animals independent of those sequenced (n = 3–4 per group): (a) *Retsat*, (b) *Enpp6*, (c) *Polq*. Bars show mean ± SEM with individual animals overlaid; open circles denote saline, filled circles nicotine. Each transcript was assessed by three-way ANOVA with sex, genotype, and treatment as factors. Significant main effects of genotype were detected for *Retsat* (*F*(1,24) = 28.32, *P* < 0.0001), *Enpp6* (*F*(1,24) = 14.12, *P* = 0.001), and *Polq* (*F*(1,24) = 9.76, *P* = 0.005). *Retsat* and *Polq* were elevated and *Enpp6* reduced in α6^GG^ animals relative to α6^CC^. No main effects of treatment or genotype-by-treatment interactions were detected. Brackets denote genotype main effects. **(d)** MDA concentration in VTA homogenates from saline- and nicotine-treated α6^CC^ and α6^GG^ animals. Bars show mean ± SEM with individual animals overlaid; open circles denote saline, filled circles nicotine. Baseline MDA was elevated in saline-treated α6^GG^ animals relative to saline-treated α6^CC^ animals (*P* = 0.04), evaluated by a one-tailed planned comparison specified a priori on the basis of the *Retsat* and *Polq* upregulation identified by transcriptomic analysis. Treatment effects within each genotype were assessed by Tukey-corrected post-hoc comparison following a restricted maximum likelihood model. Nicotine exposure significantly reduced MDA in α6^GG^ animals (*P* = 0.01) and produced no change in α6^CC^ animals (*P* = 0.96).

### Nicotine produces a genotype-dependent shift in VTA lipid peroxidation

The consistent upregulation of *Retsat* and *Polq* across every α6^GG^ group generated a specific directional prediction where the α6^GG^ VTA sustains greater oxidative load. We tested this prediction directly by quantifying malondialdehyde (MDA), a byproduct of polyunsaturated fatty acid peroxidation and an established index of tissue oxidative stress [37], in VTA homogenates (**Fig. 2d**). Baseline MDA was elevated in saline-treated α6^GG^ animals relative to α6^CC^ controls (*p* = 0.04, planned one-tailed comparison specified a priori based on the transcriptomic prediction). The α6^GG^ VTA therefore sustains a higher basal level of lipid peroxidation. Nicotine exposure, however, altered this state in a genotype-dependent manner. α6^CC^ animals showed no change in MDA between saline and nicotine conditions (*p* = 0.96), whereas nicotine exposure significantly reduced MDA in α6^GG^ animals (*p* = 0.01), bringing lipid peroxidation in this genotype to a level comparable to that of α6^CC^ animals.

Nicotine therefore produces a genotype-dependent effect on the VTA that is not detectable at the level of transcript abundance. No gene reached significance for treatment in any of the four sex-by-genotype comparisons, yet the biochemical consequence is measurable. This result also connects the components of the core signature. *Retsat* and *Enpp6* both act on membrane lipid metabolism, the latter supplying choline for phosphatidylcholine synthesis in myelinating oligodendrocytes, and MDA reports peroxidation of those same membrane lipids [33–34, 37–38]. A genotype that differs in lipid-handling enzymes at baseline, in lipid peroxidation at baseline, and in the response of that peroxidation to nicotine is one in which membrane composition is a plausible locus of vulnerability.

### The *Chrna6* co-expression network shows sex- and genotype-dependent organization

Next, we asked whether the variant alters not only which transcripts are expressed but the relationships among them. Because *Chrna6* abundance itself did not differ between genotypes, we examined the transcripts with which *Chrna6* co-varies. Transcripts correlated with *Chrna6* were identified within each sex, genotype, and treatment group (n = 9 per group), with correlations corrected across all transcripts by the Benjamini–Hochberg procedure (**Fig. 3**).

**Figure 3.**
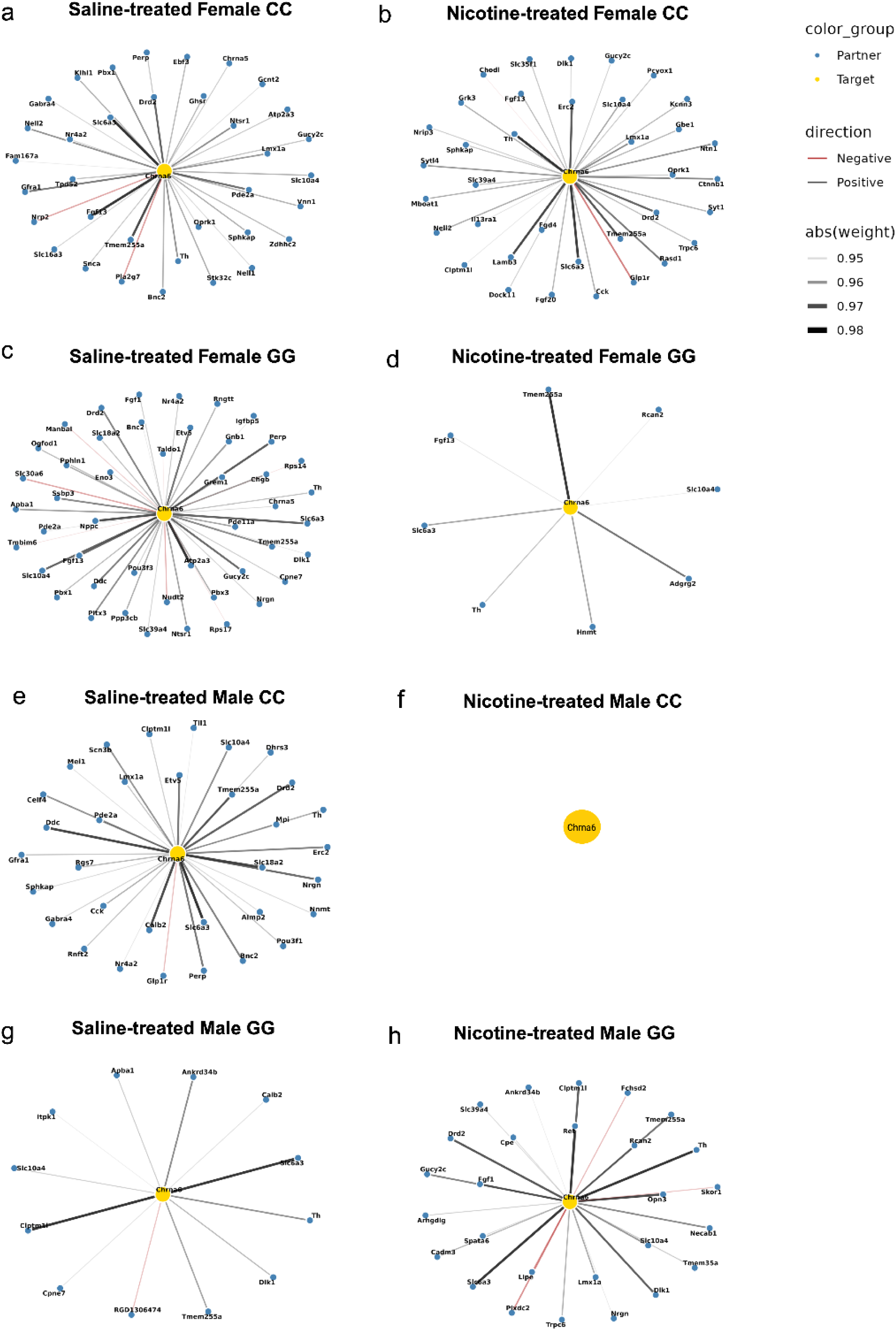
*Chrna6* co-expression networks across sex, genotype, and treatment. Gene networks showing transcripts correlated with *Chrna6* within each experimental group (n = 9 per group): **(a)** saline-treated α6^CC^ females, **(b)** nicotine-treated α6^CC^ females, **(c)** saline-treated α6^GG^ females, **(d)** nicotine-treated α6^GG^ females, **(e)** saline-treated α6^CC^ males, **(f)** nicotine-treated α6^CC^ males, **(g)** saline-treated α6^GG^ males, **(h)** nicotine-treated α6^GG^ males. *Chrna6* is shown as the central target node (yellow) and correlated transcripts as partner nodes (blue). Pearson correlations were computed within groups across all expressed transcripts and corrected by the Benjamini–Hochberg procedure at q < 0.05; only transcripts surviving correction are displayed. Edge shading indicates the absolute correlation coefficient, from 0.95 to 0.99, and edge color indicates the direction of the correlation (black, positive; red, negative). No transcripts survived correction in nicotine-treated α6^CC^ males **(f)**, which is represented by the target node alone. Dopamine synthesis and transport machinery, including *Th*, *Slc6a3*, *Drd2*, *Ddc*, and *Slc18a2*, together with the midbrain dopaminergic identity factors *Lmx1a*, *Nr4a2*, and *Pitx3*, recurs across groups. Network composition following nicotine exposure differs between genotypes in opposite directions in the two sexes.

Across nearly every group, *Chrna6* was correlated with the machinery of dopamine synthesis, packaging, and reuptake. Tyrosine hydroxylase (*Th*) and the dopamine transporter (*Slc6a3*) appeared in seven of eight networks, accompanied in most groups by the D2 autoreceptor (*Drd2*), and in several by aromatic L-amino acid decarboxylase (*Ddc*) and the vesicular monoamine transporter (*Slc18a2*). The midbrain dopaminergic identity factors *Lmx1a*, *Nr4a2*, and *Pitx3* were also present, as well as *Slc10a4*, a vesicular carrier expressed in monoaminergic and cholinergic neurons. The consistent presence of these partners across sexes, genotypes, and treatments indicates that *Chrna6* co-varies with the dopaminergic phenotype of the VTA, as expected for a receptor subunit whose expression is highly specific to this region.

Under saline conditions, three of four groups maintained extensive networks built around this dopaminergic core. Saline-treated α6^CC^ females (**Fig. 3a**) and α6^GG^ females (**Fig. 3c**) both showed broad connectivity, the latter including *Ddc*, *Slc18a2*, and *Pitx3* alongside *Th* and *Slc6a3*, as did saline-treated α6^CC^ males (**Fig. 3e**). Saline-treated α6^GG^ males were the exception, showing a relatively restricted network retaining *Th*, *Slc6a3*, and *Slc10a4* but few additional partners (**Fig. 3g**).

Nicotine exposure reorganized these relationships in a manner that depended on both sex and genotype, and in opposite directions between the sexes. In females, α6^CC^ animals retained an extensive network after nicotine, with the dopaminergic core preserved and additional partners appearing among synaptic release and neuropeptide signaling transcripts, including *Syt1*, *Erc2*, *Oprk1*, *Glp1r*, and *Cck* (**Fig. 3b**). α6^GG^ females showed the opposite response: the broad saline network contracted to a small number of transcripts, retaining *Th*, *Slc6a3*, *Slc10a4*, *Fgf13*, and *Tmem255a* while losing most of its partners (**Fig. 3d**). In males, the pattern was reversed. α6^CC^ animals, which showed extensive connectivity under saline conditions, showed no transcripts correlated with *Chrna6* after nicotine (**Fig. 3f**). α6^GG^ males, which had the most restricted network at baseline, showed the opposite shift, with connectivity expanding after nicotine to include *Th*, *Slc6a3*, *Drd2*, and *Lmx1a* alongside transcripts associated with neurotrophic signaling and structural remodeling, among them *Ret*, *Cadm3*, *Plxdc2*, *Necab1*, and *Cpe* (**Fig. 3h**). The genotype associated with reduced connectivity after nicotine therefore differs between the sexes: α6^GG^ in females, α6^CC^ in males. Within each sex, the two genotypes respond to nicotine in opposing directions, and the direction of that response is inverted between sexes.

### Tyrosine hydroxylase expression and dopaminergic neuron number follow the same sex-by-genotype pattern as gene network reorganization

The co-expression analysis identified *Th* as a partner of *Chrna6*, and the groups showing network expansion after nicotine were α6^CC^ females and α6^GG^ males. We therefore asked whether tyrosine hydroxylase itself differs between these groups at the protein level, in drug-naive animals. TH protein expression was measured by western blot in VTA punches from P31 animals (**Fig. 4a– c**). Two-way ANOVA identified a significant sex-by-genotype interaction (*F*(1,19) = 17.23, *P* = 0.0005). Female α6^CC^ animals showed higher TH expression than female α6^GG^ animals (*P* = 0.002), while male α6^GG^ animals showed higher expression than male α6^CC^ animals (*P* = 0.027). No sex or genotype differences were detected for D1, D2, D3, or DAT protein in this region (**Supplementary Fig. S1a–d**), indicating that the effect is specific to the rate-limiting enzyme of dopamine synthesis rather than reflecting a general shift in dopaminergic protein expression. In the nucleus accumbens, TH protein showed a main effect of sex, with higher expression in males regardless of genotype (*F*(1,17) = 6.06, *P* = 0.025), and no differences were detected for D2 or DAT (**Supplementary Fig. S1e–g**). The sex-by-genotype pattern is therefore specific to the VTA.

**Figure 4.**
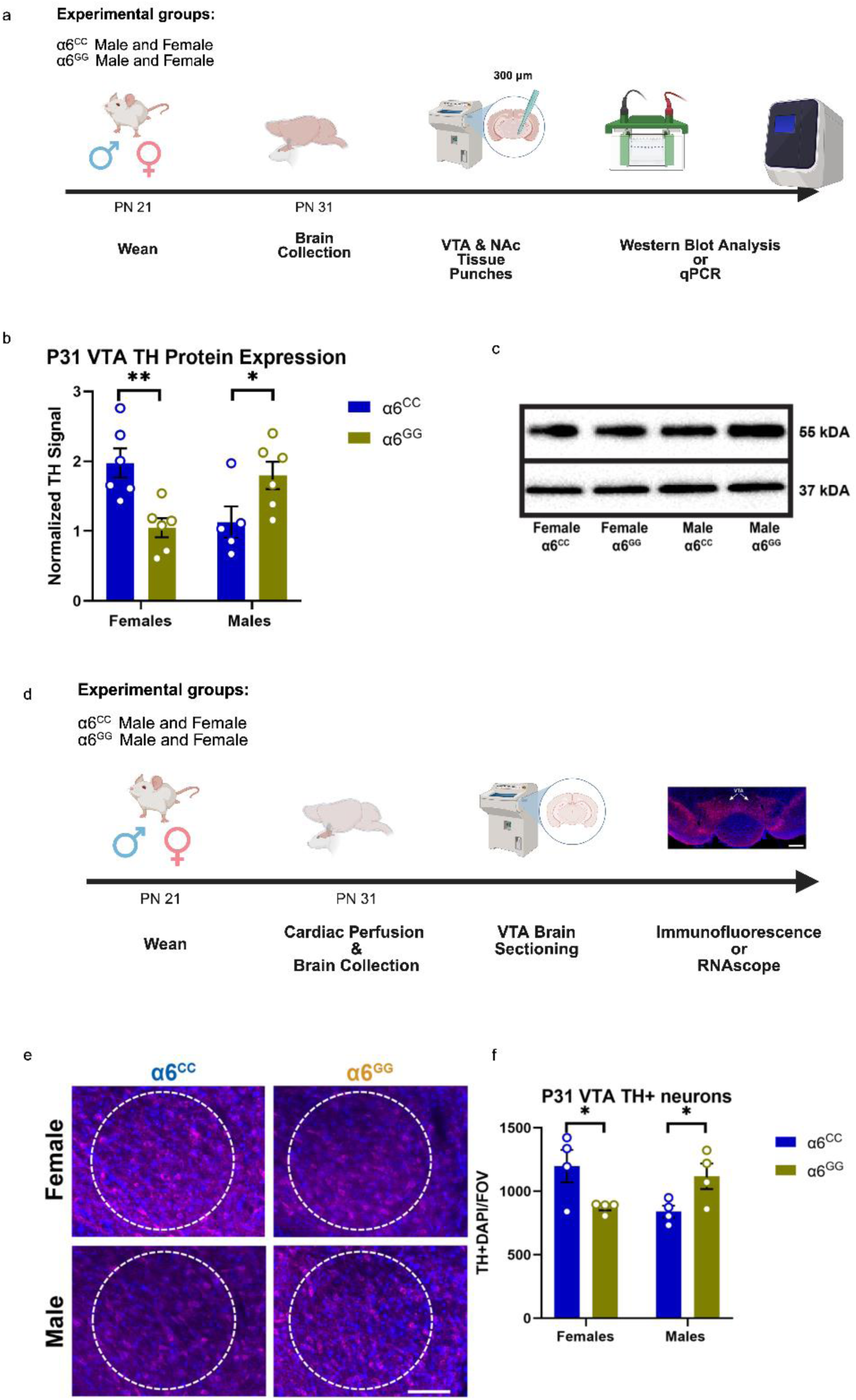
Tyrosine hydroxylase expression and dopaminergic neuron number in the adolescent VTA follow a sex-by-genotype pattern. **(a)** Experimental timeline for western blot and qPCR analyses. Male and female α6^CC^ and α6^GG^ rats were weaned at PN 21 and brains collected at PN 31. Tissue was sectioned at 300 µm and the VTA and NAc isolated by punch. **(b)** TH protein expression in the VTA measured by western blot. Two-way ANOVA identified a significant sex-by-genotype interaction (*F*(1,19) = 17.23, *P* = 0.0005). Female α6^CC^ animals showed higher TH expression than female α6^GG^ animals (\*\**P* = 0.002), while male α6^GG^ animals showed higher expression than male α6^CC^ animals (\**P* = 0.027). n = 5–6 per group. **(c)** Representative western blot images. TH, 55 kDa; GAPDH loading control, 37 kDa. **(d)** Experimental timeline for immunofluorescence and RNAscope analyses. Animals were weaned at PN 21 and underwent cardiac perfusion and brain collection at PN 31, followed by VTA sectioning. Inset shows a representative coronal section of the VTA (4× magnification). Scale bar, 250 µm. **(e)** Representative images (20× magnification) of TH (magenta) and DAPI (blue) staining in the VTA of female and male α6^CC^ and α6^GG^ animals. Dashed circles indicate the region of interest. Scale bar, 150 µm. **(f)** Quantification of TH-positive neurons. Two-way ANOVA identified a significant sex-by-genotype interaction (*F*(1,12) = 12.53, *P* = 0.004). Female α6^CC^ animals showed more TH-positive neurons than female α6^GG^ animals (\**P* = 0.019), while male α6^GG^ animals showed more than male α6^CC^ animals (\**P* = 0.04). n = 4 per group. Animals were drug-naive at the time of collection. Bars show mean ± SEM with individual animals overlaid; data points represent independent biological samples. \**P* < 0.05, \*\**P* < 0.01.

To determine whether the elevated TH signal reflected greater synthesis capacity per neuron or a greater number of dopaminergic neurons, we quantified TH-positive cells by immunofluorescence, identifying dopaminergic neurons by colocalization of TH and DAPI (**Fig. 4d–f**). Two-way ANOVA identified a significant sex-by-genotype interaction (*F*(1,12) = 12.53, *P*= 0.004). Female α6^CC^ animals showed more TH-positive neurons than female α6^GG^ animals (*P* = 0.019), and male α6^GG^ animals showed more than male α6^CC^ animals (*P* = 0.04). The difference therefore involves dopaminergic neuron number rather than TH expression per cell. These animals were drug-naive at the time of collection, so the sex-by-genotype difference in dopaminergic capacity is present before any nicotine exposure. The groups carrying elevated TH and greater dopaminergic neuron number, α6^CC^ females and α6^GG^ males, are the same groups whose *Chrna6* co-expression networks expand following nicotine, and the same groups in which sub-chronic nicotine produces increased locomotor and anxiolytic responses. Two independent measures, obtained in different animals by different methods, converge on the same sex-by-genotype pattern, suggesting that the genotype associated with greater dopaminergic capacity differs between the sexes.

### The risk allele alters predicted 3′ UTR conformation

Having established that the variant reshapes the molecular and biochemical environment of the VTA without altering *Chrna6* abundance, we turned to the question of mechanism. Because rs2304297 lies within a non-coding regulatory region, we tested the routes through which a 3′ UTR variant can act. *In silico* folding of the *CHRNA6* 3′ UTR carrying either allele (C or G) predicted substantial structural divergence. Mountain plots derived from minimum free energy, partition function, and centroid structures differed between alleles, with the largest deviations at and downstream of the polymorphic site at position 123 (**Fig. 5a-b, e-f**). Positional entropy, which quantifies thermodynamic uncertainty at each nucleotide, was higher and more frequently elevated around and downstream of the SNP in the G allele, indicating greater conformational flexibility (the C allele showed lower entropy and a more thermodynamically constrained local conformation) (**Fig. 5c-d**). Because 3′ UTR folding governs the accessibility of regulatory motifs to the machinery that recognizes them, these predictions provided a testable rationale for genotype-dependent differences in transcript processing.

**Figure 5.**
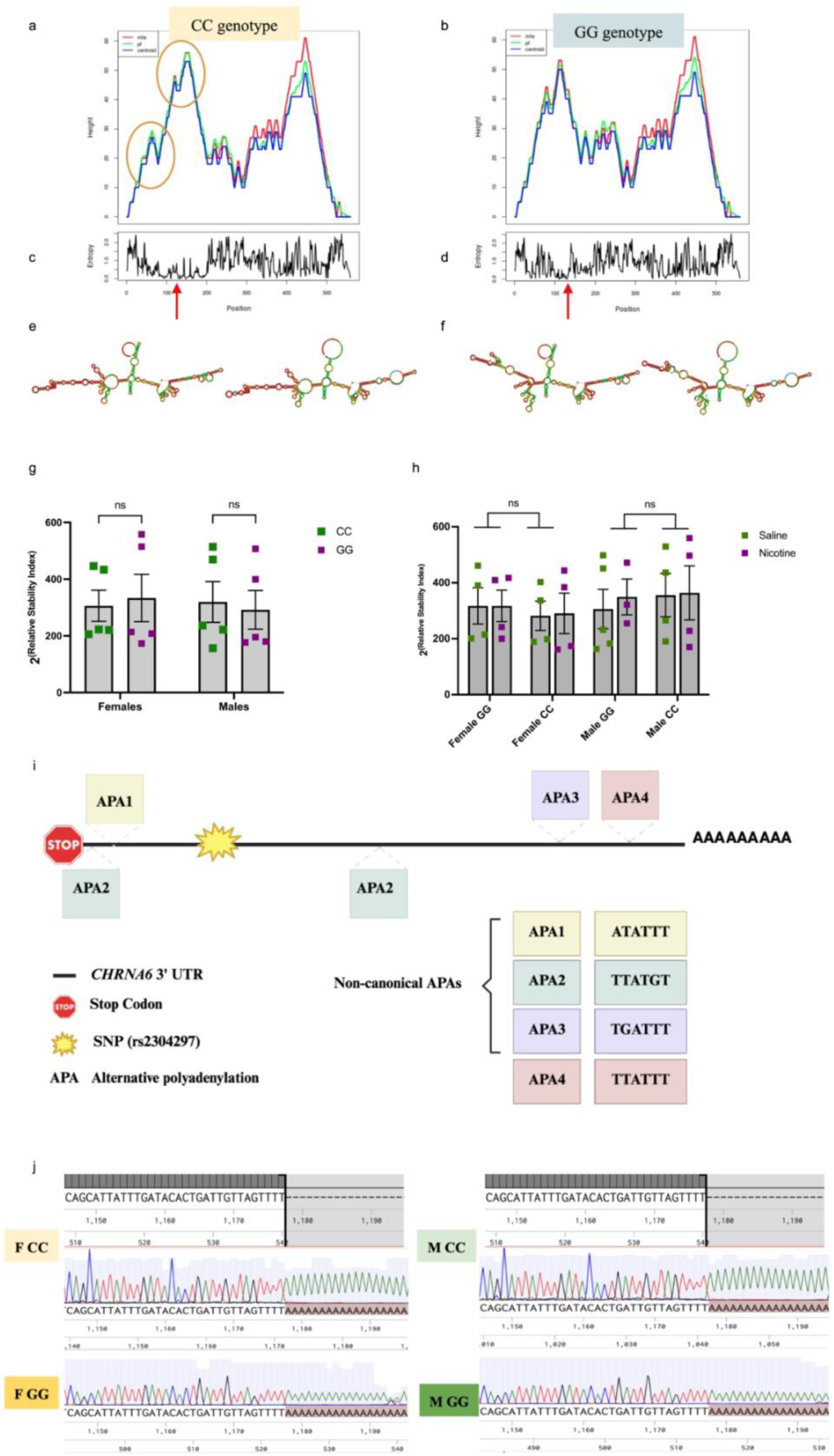
The rs2304297 allele alters predicted 3′ UTR conformation without changing transcript stability or polyadenylation. **(a, b)** Mountain plot representations of the *CHRNA6* 3′ UTR carrying the **(a)** C allele and **(b)** G allele, generated in ViennaRNA. Each plot shows minimum free energy (red), partition function (green), and centroid (blue) structures. Circled regions mark positions of divergence between alleles. **(c, d)** Positional entropy across the 3′ UTR for the **(C)** C allele and **(d)** G allele. Higher values indicate greater thermodynamic uncertainty and conformational flexibility at a given nucleotide. Red arrows mark the polymorphic site at position 123. The G allele shows higher and more frequent entropy peaks at and downstream of the SNP. **(e, f)** Predicted secondary structures for the **(E)** C allele and **(F)** G allele, colored by base-pairing probability. **(g)** Transcript stability at baseline, expressed as 2^-(relative^ ^stability^ ^index)^, calculated as mature *Chrna6* mRNA relative to unspliced pre-mRNA. Green denotes α6^CC^ and purple α6^GG^ (N = 5). Two-way ANOVA identified no effect of genotype (F(1,15) = 1.26, *P* = 0.280), sex (F(1,15) = 0.76, *P* = 0.397), or their interaction. **(h)** Transcript stability across treatment conditions (N = 3–5 per group). Green denotes saline and purple nicotine. Three-way ANOVA identified no effect of treatment (F(1,24) = 0.08, *P* = 0.776), genotype (F(1,24) = 0.03, *P* = 0.867), sex (F(1,24) = 0.46, *P* = 0.503), or any interaction, including genotype-by-treatment (F(1,24) = 0.15, *P* = 0.699). ns, not significant. **(i)** Schematic of the humanized *CHRNA6* 3′ UTR showing the stop codon, the rs2304297 polymorphic site (star), and four candidate alternative polyadenylation motifs: APA1 (ATATTT), APA2 (TTATGT, present at two loci), APA3 (TGATTT), and APA4 (TTATTT), positioned both upstream and downstream of the SNP. **(j)** Representative Sanger sequencing traces from 3′ RACE amplicons cloned from female α6^CC^, female α6^GG^, male α6^CC^, and male α6^GG^ animals. A single conserved polyadenylation site was identified across all groups, with no genotype- or sex-dependent difference in APA site usage.

### The rs2304297 variant does not alter *Chrna6* transcript stability

We then tested whether the predicted structural difference alters transcript turnover, quantifying mature *Chrna6* mRNA relative to unspliced pre-mRNA using assays specific to the spliced exon–exon junction and to the intron 5/exon 6 junction respectively. At baseline, two-way ANOVA identified no effect of genotype (F(1,15) = 1.26, *P* = 0.28), sex (F(1,15) = 0.76, *P* = 0.39), or their interaction (F(1,15) = 1.26, *P* = 0.28) (**Fig. 5g**). Because nicotine can engage 3′ UTR-dependent stabilization pathways, and because an effect might be revealed only under drug challenge, we repeated the measurement under nicotine treatment. Three-way ANOVA identified no effect of treatment (F(1,24) = 0.08, *P* = 0.776), genotype (F(1,24) = 0.03, *P* = 0.867), or sex (F(1,24) = 0.46, *P* = 0.503), and no interactions, including the genotype-by-treatment interaction of primary interest (F(1,24) = 0.15, *P* = 0.699) (**Fig. 5h**). Altered transcript degradation is therefore excluded as a mechanism for this variant, under both saline- and nicotine-treated conditions.

### The variant does not alter polyadenylation site selection

We next tested whether the predicted conformational change alters which polyadenylation signal the cleavage machinery uses, a mechanism that would produce isoforms differing in 3′ UTR length and therefore in regulatory content. Four candidate polyadenylation motifs were identified within the humanized 3′ UTR, positioned both upstream and downstream of the SNP: APA1 (ATATTT), APA2 (TTATGT, occurring at two distinct loci), APA3 (TGATTT), and APA4 (TTATTT) (**Fig. 5i**). We characterized isoform usage *in vivo* by 3′ Rapid Amplification of cDNA Ends (RACE), using a gene-specific primer positioned upstream of the first predicted site so that both canonical and alternative isoforms would amplify. Amplicons were cloned and sequenced bidirectionally across all four sex-and-genotype groups. Sequencing revealed a single conserved polyadenylation site, with no detectable genotype- or sex-dependent difference in site usage (**Fig. 5j**).

Taken together, **Figures 5g-j** exclude the two mechanisms most invoked for 3′ UTR variants. rs2304297 alters predicted conformation but changes neither transcript turnover nor isoform length. Its regulatory effect must therefore operate at the level of RNA–protein or RNA– RNA interaction, consistent with the reduced local translation signature already observed in α6^GG^ females.

### miRNA-365-5p engagement with *Chrna6* is genotype-dependent

Among microRNAs predicted to target the human *CHRNA6* 3′ UTR, miRNA-365-5p is conserved in rats and humans and is predicted to bind at positions 93–99, adjacent to but not overlapping the polymorphic site. Because the predicted conformational change extends across the 3′ UTR rather than being confined to position 123, a variant at that position could plausibly alter accessibility at this non-overlapping locus.

Correlation of individual ΔCt values across biological replicates revealed a relationship dependent on both sex and genotype (**Fig. 6 a-d**). In females, a significant positive correlation between miRNA-365-5p and *Chrna6* was present in α6^GG^ animals (R² = 0.556, *P* = 0.008) and absent in α6^CC^ animals (*P* = 0.538). The pattern inverted in males, with a significant positive correlation in α6^CC^ animals (R² = 0.339, *P* = 0.045) and none in α6^GG^ animals (*P* = 0.538). A positive relationship between a microRNA and its predicted target is not the inverse relationship expected of canonical degradation, consistent with the stability data above, which showed no genotype effect on transcript turnover.

**Figure 6.**
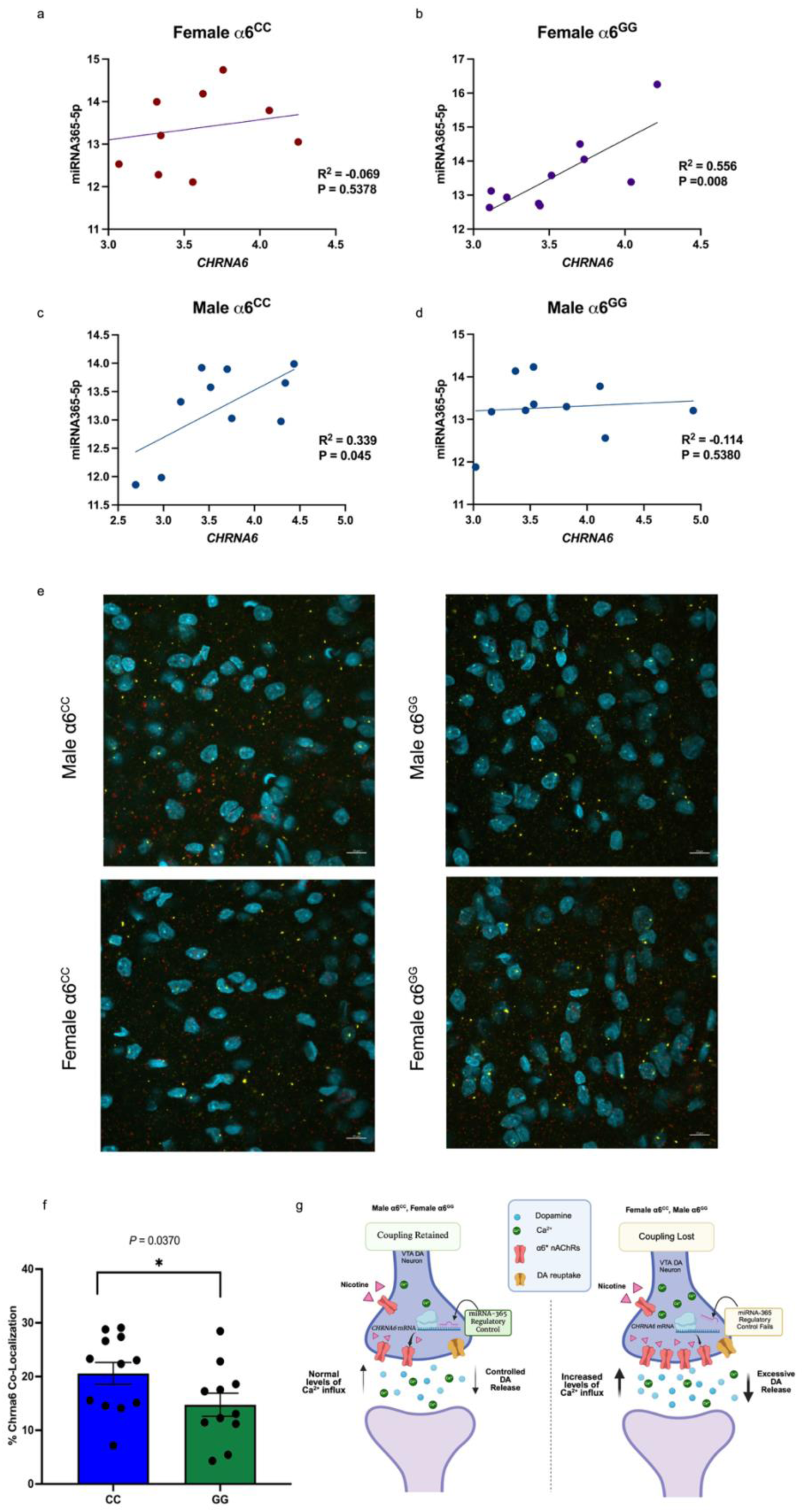
miRNA-365-5p engagement with *Chrna6* differs by genotype and sex, and a proposed model of its consequences. **(a-d)** Correlation between miRNA-365-5p and *Chrna6* expression, plotted as individual ΔCt values across biological replicates **(a)** female α6^CC^, **(b)** female α6^GG^, **(c)** male α6^CC^, **(d)** male α6^GG^. A significant positive correlation was present in α6^GG^ females (*P* = 0.008) and α6^CC^ males (*P* = 0.045), and absent in α6^CC^ females (*P* = 0.538) and α6^GG^ males (*P* = 0.538). Lines show linear fits. **(e)** Representative multiplex fluorescent *in situ* hybridization in the posterior VTA of male α6^CC^, male α6^GG^, female α6^CC^, and female α6^GG^ animals. Cyan, DAPI-labelled nuclei; yellow, *Chrna6*; red, miRNA-365-5p. Images acquired at 63× magnification. Scale bar, 10 µm. **(f)** Quantification of *Chrna6*–miRNA-365-5p colocalization by genotype, measured by an object-based centre-of-mass algorithm across five fields of view per section. Bars show mean ± SEM with individual animals overlaid. A three-way ANOVA incorporating sex, genotype identified a significant main effect of genotype (F(1,15) = 5.24, \**P* = 0.037), with reduced colocalization in α6^GG^ animals. The omnibus model did not reach significance (F(7,15) = 1.63, *P* = 0.203). **(g)** Proposed model. Reduced *Chrna6*–miRNA-365-5p colocalization and elevated basal lipid peroxidation are genotype-driven and present in α6^GG^ animals of both sexes. Whether reduced colocalization results in loss of functional coupling between the two transcripts depends on sex, and does so in opposite directions in males and females. **Left**, coupling retained (male α6^CC^, female α6^GG^): miRNA-365-5p maintains translational constraint on α6*nAChR density, nicotine produces controlled calcium influx and stable dopamine release, tyrosine hydroxylase expression and dopaminergic neuron number remain at baseline, the *Chrna6* co-expression network does not expand following nicotine, and these groups are behaviorally unresponsive to sub-chronic nicotine. **Right**, coupling lost (female α6^CC^, male α6^GG^): translational constraint is released, predicting elevated α6*nAChR density, increased calcium influx, and excessive dopamine release; these groups show elevated tyrosine hydroxylase expression and dopaminergic neuron number, expansion of the co-expression network following nicotine, and the behavioral responses to sub-chronic nicotine previously reported in these lines. α6*nAChR density, calcium influx, and dopamine release magnitude are proposed rather than measured.

A regulatory interaction requires the two transcripts to be present in the same cells. We used multiplex fluorescent *in situ* hybridization to measure how frequently *Chrna6* and miRNA-365-5p colocalize within individual cells of the VTA (**Fig. 6e**). Colocalization was quantified by an object-based center-of-mass algorithm across five fields of view per section. A three-way ANOVA incorporating sex, genotype, and anatomical subregion identified a significant main effect of genotype (F(1,15) = 5.24, p = 0.037), with reduced *Chrna6*–miRNA-365-5p colocalization in α6^GG^ animals (**Fig. 6f**). A three-way ANOVA incorporating sex, genotype identified a significant main effect of genotype (*F*(1,15) = 5.24, *P* = 0.037), with reduced *Chrna6*– miRNA-365-5p colocalization in α6^GG^ animals (Fig. 5f). No other main effect or interaction reached significance, and the omnibus model did not reach significance (*F*(7,15) = 1.63, *P* = 0.203).

The rs2304297 genotype is therefore associated with reduced spatial engagement of miRNA-365-5p with the *Chrna6* transcript, in the absence of any detectable effect on transcript abundance, stability, or polyadenylation. Across these experiments, a consistent pattern emerges. The rs2304297 variant does not change how much *Chrna6* transcript is present, how quickly it turns over, or which isoform is produced. What it changes is the molecular context in which that transcript operates. A signature spanning myelination, lipid metabolism, and DNA repair distinguishes the two genotypes in both sexes. The α6^GG^ VTA sustains elevated basal lipid peroxidation, and nicotine normalizes this measure in α6^GG^ animals while leaving α6^CC^ animals unchanged. Engagement of a conserved regulatory microRNA with the transcript is reduced in α6^GG^ animals. The variant acts on the environment surrounding *Chrna6*, not on *Chrna6* abundance itself.

Taken together, the rs2304297 variant establishes a distinct molecular environment in the adolescent VTA. A reproducible signature spanning myelin maintenance, lipid metabolism, and genomic repair distinguishes the genotypes in both sexes. The α6^GG^ VTA maintains elevated lipid peroxidation, confirming biochemically what the transcriptional signature predicted. Spatial engagement of miRNA-365-5p with the *Chrna6* transcript is reduced in α6^GG^ animals. However, none of this involves the transcript’s stability or polyadenylation site selection.

## Discussion

Humanized rats carrying the C or G allele of the human *CHRNA6* 3′ untranslated region show no baseline behavioral differences. However, they respond to adolescent nicotine in a sex- and genotype-dependent manner, with α6^CC^ females and α6^GG^ males showing enhanced locomotor and anxiolytic responses [20], enhanced nicotine plus cue-primed reinstatement [39], and altered dopamine overflow in the nucleus accumbens shell [21]. The mechanism by which a single base change outside the coding sequence causes this pattern has remained unknown. Cardenas et al. [20] reported that this SNP does not alter α6 subunit mRNA expression in the ventral tegmental area (VTA), substantia nigra, or interpeduncular nucleus, and proposed that the effect may instead involve the regulatory functions of the 3′ untranslated region, specifically interaction with RNA-binding proteins or microRNAs. Here we examined the molecular consequences of rs2304297 *in vivo* to uncover its downstream impact.

Our findings suggest that this variant does not alter *Chrna6* transcript abundance, turnover, or polyadenylation site selection, but it does establish a distinct molecular environment in the adolescent VTA and reduces spatial interaction of miRNA-365-5p with the *Chrna6* transcript. The consequences of these genotype-dependent effects, however, depend on sex. Regulatory coupling between miRNA-365-5p and *Chrna6* is lost in α6^CC^ females and α6^GG^ males, which are the groups that show elevated dopaminergic function and behavioral sensitivity to nicotine in this model.

### rs2304297 does not act through transcript abundance, turnover, or isoform selection

The regulatory functions of 3′ untranslated regions are well characterized. They determine transcript stability, subcellular localization, translational efficiency, and susceptibility to microRNA-mediated repression [40–41]. Variants within this region impact human disease through several of these routes. For instance, polyadenylation site selection is altered by rs10954213 in the *IRF5* 3′ UTR, which strengthens a proximal signal and shortens the transcript, increasing protein output and elevating autoimmune risk [23]. Moreover, the rs12720208 variant in the *FGF20* 3′ UTR disrupts a miRNA-433 binding site, releasing translational repression and driving α-synuclein pathology in Parkinson’s disease [24]. Within the mesolimbic system, the variable number tandem repeat in the *SLC6A3* 3′ UTR alters dopamine transporter density and constitutes an established risk factor for substance use disorders [42].

The humanized rat model allows these mechanisms to be tested in tissue. We therefore assessed transcript stability and polyadenylation site selection and evaluated microRNA engagement as an index of the remaining routes. Transcript stability, measured as mature *Chrna6* relative to unspliced pre-mRNA, did not differ by genotype at baseline or with nicotine exposure. Polyadenylation site selection, assessed by 3′ RACE and sequencing across all four sex-and-genotype groups, revealed a single conserved site. In addition, *Chrna6* itself was not differentially expressed between genotypes in any comparison, consistent with the absence of α6 mRNA differences reported by *in situ* hybridization in this model [20]. Excluding these routes in tissue leaves only mechanisms that preserve transcript quantity and structure, acting instead at the level of translation or localization.

### The SNP alters the conformational ensemble of the 3′ untranslated region

Computational folding predicted that the G allele adopts a more conformationally flexible structure, with elevated positional entropy at and downstream of the polymorphic site. A single nucleotide variant that alters the conformational ensemble of a transcript without changing the presence of its regulatory motifs has been identified as a riboSNitch [43], which are concentrated in untranslated regions [44]. Because 3′ UTR folding governs the accessibility of regulatory motifs to the machinery that recognizes them, the conformational and entropy differences we observe provide a testable rationale for genotype-dependent differences in transcript handling without any change in the motifs themselves.

### The α6^GG^ VTA sustains a distinct metabolic and redox state

Three transcripts differed by genotype in every sex-and-treatment comparison and replicated in an independent cohort of animals. *Enpp6*, a choline-specific phosphodiesterase required for oligodendrocyte differentiation and myelin membrane synthesis [33], was reduced in α6^GG^ rats, while *Retsat*, encoding retinol saturase [34], and *Polq*, encoding the polymerase that resolves double-strand breaks by microhomology-mediated end joining [35–36], were elevated. Two of these genes act on membrane phospholipid metabolism and the third responds to the genomic damage that oxidative stress produces. The pattern appeared in saline-treated animals of both sexes, so it describes how the α6^GG^ VTA differs before any drug exposure. We tested the prediction that follows from it. Basal lipid peroxidation was elevated in α6^GG^ animals, establishing that the expression difference reflects a real difference in oxidative state rather than one confined to transcript levels. The relevance of this state to dopaminergic neurons specifically is well established. Midbrain dopaminergic neurons are vulnerable to oxidative stress, in part because their activity imposes a sustained calcium load on mitochondria that generates reactive oxygen species [45–46].

The reduced *Enpp6* effect is of particular interest given the developmental window. Myelination of the mesolimbic pathway continues through adolescence [47–48], and limited choline availability for phosphatidylcholine synthesis would reduce the biosynthetic capacity supporting that process [49]. Such a difference would not produce an acute behavioral phenotype, consistent with the absence of baseline behavioral differences in this model across food reinforcement, locomotor activity, and three measures of anxiety-like behavior [20], but could impact how the circuit responds to nicotine during the same period.

### Sex and genotype shape how the *Chrna6* network responds to nicotine

Across nearly every experimental group, *Chrna6* was correlated with the machinery of dopamine synthesis, packaging, and reuptake, including *Th*, *Slc6a3*, *Drd2*, *Ddc*, and *Slc18a2*, together with the midbrain identity factors *Lmx1a* and *Nr4a2*. This is the expected result for a subunit whose expression is highly concentrated in the catecholaminergic nuclei [50, 8, 51]. What differed between groups was how this network responded to nicotine. In females, α6^CC^ animals maintained an extensive network and recruited synaptic release and neuropeptide signaling transcripts, while α6^GG^ animals lost most of their baseline connectivity. In males the pattern reversed, with α6^CC^ animals losing all detectable connectivity and α6^GG^ animals expanding a restricted baseline network to include dopaminergic and neurotrophic transcripts. These analyses were conducted at nine animals per group and are descriptive, but the direction of the pattern is notable. The two groups whose networks expand following nicotine, α6^CC^ females and α6^GG^ males, are the two groups in which this model shows enhanced nicotine-induced locomotor and anxiolytic behavior [20], enhanced nicotine plus cue-primed reinstatement [39], and enhanced drug-induced dopamine overflow in the nucleus accumbens shell [21].

The protein data divide the groups the same way. In drug-naive animals, tyrosine hydroxylase expression and dopaminergic neuron number in the VTA were elevated in α6^CC^ females and α6^GG^ males relative to their counterparts [52]. The effect appears specific because no genotype differences were detected for D1, D2, D3, or DAT protein, and in the nucleus accumbens TH varied with sex alone. These observations align with the neurochemical signature established in this model. Drug-naive adolescent α6^GG^ males show elevated tissue dopamine in the medial prefrontal cortex and nucleus accumbens core relative to α6^CC^ males [53], and adolescent α6^GG^ males show enhanced methamphetamine-induced dopamine overflow in the nucleus accumbens shell, while α6^CC^ females show enhanced nicotine- and methamphetamine-induced overflow relative to α6^GG^ females [21]. Increased dopaminergic capacity in α6^CC^ females and α6^GG^ males is therefore supported by tissue-level neurotransmitter content, increased release, synthetic enzyme expression, and dopaminergic neuron numbers measured across separate studies.

### A microRNA interaction that does not degrade the transcript

MicroRNAs do not only act through transcript degradation. Argonaute-loaded silencing complexes localize to cytoplasmic processing bodies, and transcripts repressed at the level of translation accumulate there in a microRNA-dependent manner [54–55]. Therefore, transcripts are stored rather than degraded and can be released when needed [56], so that microRNA binding reduces protein output while leaving transcript abundance intact [57]. For instance, the rs3800373 polymorphism in the *FKBP5* 3′ UTR alters local secondary structure and reduces miRNA-320a binding efficiency, increasing its translation without a change in *FKBP5* mRNA levels. This difference mediates vulnerability to chronic post-traumatic pain [58]. We propose that rs2304297 works the same way, altering how readily miRNA-365-5p engages with *Chrna6* transcript.

miRNA-365-5p is conserved between rat and human and is predicted to bind the *CHRNA6* 3′ UTR at positions 93–99, adjacent to the polymorphic site at position 123. Because the predicted conformational difference extends across the untranslated region, a variant at position 123 could alter accessibility at this locus. Cardenas et al. [20], in proposing microRNA interaction as the likely mechanism for this variant, cited the observation that a functional 3′ UTR polymorphism in prodynorphin (rs2235749) alters microRNA-365 binding in ventral striatonigral neurons and influences novelty seeking and positive reward traits [59]. We tested that proposal directly and found altered miRNA-365-5p engagement in α6^GG^ animals. We measured both the expression relationship between the two transcripts and their spatial overlap. Expression of miRNA-365-5p correlated positively with *Chrna6* in a sex- and genotype-dependent manner, and spatial colocalization of the two transcripts was reduced in α6^GG^ animals of both sexes. The positive direction of the correlation is unexpected, since canonical microRNA activity destabilizes the target and would produce an inverse relationship. However, the stability data resolve this. Because transcript turnover does not differ by genotype, no inverse relationship should be expected. Therefore, as described above, repression without decay produces exactly this pattern.

Consequently, we propose translational repression as the primary hypothesis. It accounts for the absence of both stability and abundance differences. It also makes a specific prediction that α6 protein should differ between genotypes in the absence of any difference in *Chrna6* mRNA levels. Testing this directly is currently limited by the absence of specific antibodies against the α6 subunit [60], a limitation noted in this model. Moreover, levels of two transcripts can increase and decrease together for a simpler reason: both are driven by the same transcription factor. However, this does not fit our data well. A variant in the *CHRNA6* 3′ UTR cannot change a transcription factor program that also controls the microRNA, and shared transcriptional regulators would not change how often the two transcripts appear in the same cells.

### Genotype determines spatial engagement while sex determines functional coupling

Colocalization and coordinated expression did not track together. Colocalization followed genotype, reduced in α6^GG^ animals of both sexes. Coordinated expression followed sex and genotype together, present in α6^CC^ males and α6^GG^ females. The two measures therefore align in males but not in females, where α6^CC^ animals show colocalization without correlation and α6^GG^ animals show correlation without colocalization. Spatial co-occurrence is therefore necessary but not sufficient for a coordinated regulatory relationship, and what determines whether co-occurrence produces coordination differs between the sexes. We propose that the variant acts through two separable layers. Genotype sets the physical opportunity for interaction by altering 3′ UTR accessibility, and this occurs in both sexes. However, whether that opportunity translates into regulation depends on sex. Colocalization and basal lipid peroxidation follow genotype, while coordinated expression, tyrosine hydroxylase protein, dopaminergic neuron number, and the reorganization of the co-expression network following nicotine each follow sex and genotype together (**Table 1**). α6^CC^ females and α6^GG^ males separate from the other two groups on every one of these measures, and they are the groups in which sub-chronic nicotine produces behavioral responses in this model.

**Table 1.** Genotype- and sex-dependent effects of rs2304297 in the adolescent VTA. Summary of measured outcomes across the four sex-and-genotype groups. Spatial colocalization of Chrna6 with miRNA-365-5p follows genotype, with reduced colocalization in α6^GG^ animals of both sexes. Coordinated expression of the two transcripts, tyrosine hydroxylase protein, dopaminergic neuron number, and the response of the Chrna6 co-expression network to nicotine each follow a sex-by-genotype pattern, in which α6^CC^ females and α6^GG^ males differ from α6^CC^ males and α6^GG^ females. Correlation values are Pearson coefficients from qPCR ΔCt measurements (Fig. 6A–D). Colocalization is from multiplex fluorescent in situ hybridization (Fig. 6E–F). Tyrosine hydroxylase protein and dopaminergic neuron number were compared between genotypes within each sex. Network composition refers to transcripts correlated with Chrna6 following nicotine exposure relative to saline. Behavioral responses to sub-chronic nicotine are as previously reported in these lines (Cardenas et al. 2022).

| | Male $\alpha 6^{CC}$ | Male $\alpha 6^{GG}$ | Female $\alpha 6^{CC}$ | Female $\alpha 6^{GG}$ |
| --- | --- | --- | --- | --- |
| Colocalization (measured) | High | Low | High | Low |
| miR-365/*Chrna6* correlation (measured) | Present | Absent | Absent | Present |
| TH protein, neuron number (measured) | Lower | Elevated | Elevated | Lower |
| Network after nicotine (measured) | No partners | Expands | Expands | Contracts |
| Baseline MDA (measured) | Lower | Elevated | Lower | Elevated |
| Behavioral response (Published) | No | Yes | Yes | No |
| $\alpha 6$ receptor density (Predicted) | Normal | Elevated | Elevated | Normal |
| Vulnerability | Lower | Higher | Higher | Lower |

What produces the sex dependence is not established here. Several possibilities are testable. miRNA-365-5p availability, or the abundance of competing targets that sequester it, may differ between sexes such that equivalent colocalization produces different degrees of occupancy. Sex differences in RNA-binding protein expression [61] could alter site accessibility independently of the variant. Each predicts a measurable difference in microRNA availability or occupancy rather than in spatial distribution.

### Why a non-coding variant matters

Most disease-associated variants identified by genome-wide association lie outside protein-coding sequence [62], and for 3′ UTR variants function is usually inferred from reporter assays in cell lines, outside the transcript’s native context. RiboSNitches are especially difficult to detect this way, since the variant alters structure rather than sequence [43]. By replacing the endogenous rat 3′ UTR with the human sequence carrying either allele, this model preserves the variant in its native genomic position and allows these mechanisms to be tested in the tissue where the receptor is expressed.

This has implications beyond *CHRNA6*. Every α6^GG^ animal shows reduced colocalization, but only α6^GG^ males show the dopaminergic and behavioral phenotype, so measuring the molecular effect alone would not identify which carriers are affected. The same logic applies to association studies. If a variant increases risk in one sex and decreases it in the other, the two effects cancel when sexes are analyzed together [63]. The human data are consistent with this. Although the G allele is generally treated as the risk variant for adolescent smoking and nicotine dependence [16–19], sex-stratified analysis has shown that rs2304297 influences smoking cessation outcome in men and not in women (adjusted *P* = 0.029 and 0.96 respectively), and no variant reaching significance within one sex stratum in that study reached significance within the other [64]. If the pattern described here holds in humans, the C allele should carry risk in females rather than being neutral, and sex-stratified analysis of rs2304297 would show it.

### Limitations

Some limitations apply to these interpretations. The transcriptomic analysis used bulk VTA tissue, which contains dopaminergic, GABAergic, and glutamatergic neurons alongside oligodendrocytes, astrocytes, and microglia. *Chrna6* expression is largely restricted to dopaminergic neurons, while the genes carrying the strongest genotype signature act principally in oligodendrocytes, so we cannot determine whether the signature reflects changes within a cell type or differences in cellular composition. Single-nucleus sequencing would resolve this.

Malondialdehyde is a bulk tissue measure and cannot identify which membrane compartment or cell type underwent peroxidation, so the connection to myelin membrane remains an inference from gene expression.

The animals were homozygous for each allele, whereas human carriers are frequently heterozygous, and whether these effects scale with allele dosage is unknown.

## Conclusion

A polymorphism in the 3′ untranslated region of *CHRNA6* is associated with adolescent smoking behavior in humans but does not alter the protein sequence. The results reported here indicate that the variant does not change how much *Chrna6* transcript is present, how rapidly it turns over, or which isoform is produced. It is instead associated with a reproducible molecular signature spanning myelin maintenance, lipid metabolism, and genomic repair, with elevated basal lipid peroxidation, and with reduced spatial engagement of a conserved regulatory microRNA whose interaction with the transcript does not consume it.

These molecular effects occur in both sexes. However, coordinated regulation between miRNA-365-5p and *Chrna6* is lost in α6^CC^ females and α6^GG^ males, and it is these groups that show elevated dopaminergic function and the nicotine-induced behavioral phenotypes documented across this model. The variant therefore establishes a condition in every animal that carries it, while sex determines whether that condition becomes consequential.

## Author Contributions

All authors contributed to the study conception and design. Material preparation, data collection and analysis were performed by Yasamin Hajy Heydary, Kaylie Shea Gurholt and Emily Marie Castro. Colony maintenance, breeding and genotyping of the humanized lines were performed by Nikki Le. The first draft of the manuscript was written by Yasamin Hajy Heydary and all authors commented on previous versions of the manuscript. All authors read and approved the final manuscript.

## Data Availability

The datasets generated and analyzed during the current study are not publicly available as they form part of ongoing research in the laboratory but are available from the corresponding author on reasonable request.

## Acknowledgements

The authors thank Nicole Renee Sparks and Madeline Karen Marie Vera-Colon for their expertise and technical guidance with the RNA sequencing analysis.

This work utilized resources of the UCI Genomics Research and Technology Hub (GRT Hub), parts of which are supported by NIH grants to the Comprehensive Cancer Center (P30CA-062203) and the UCI Skin Biology Resource Based Center (P30AR075047) at the University of California, Irvine, as well as to the GRT Hub for instrumentation (1S10OD010794-01 and 1S10OD021718-01).

This study was made possible in part through access to the Optical Biology Core Facility of the Developmental Biology Center, a shared resource supported by the Cancer Center Support Grant (CA-62203) and NIH-S10OD032327-01.

## Acknowledgments

Tobacco-Related Disease Research Program (TRDRP) award [T31IP1427C, T33IP617, T34IR8049] (S.L.); University of California Irvine (UCI) National Institute of Drugs and Abuse (NIDA) T32 training grant (T32DA050558) (EMC).

