## Supplementary Figures and Tables for "“A human *CHRNA6* 3′ UTR variant alters the molecular environment and microRNA engagement of the adolescent VTA without changing *Chrna6* transcript abundance”"

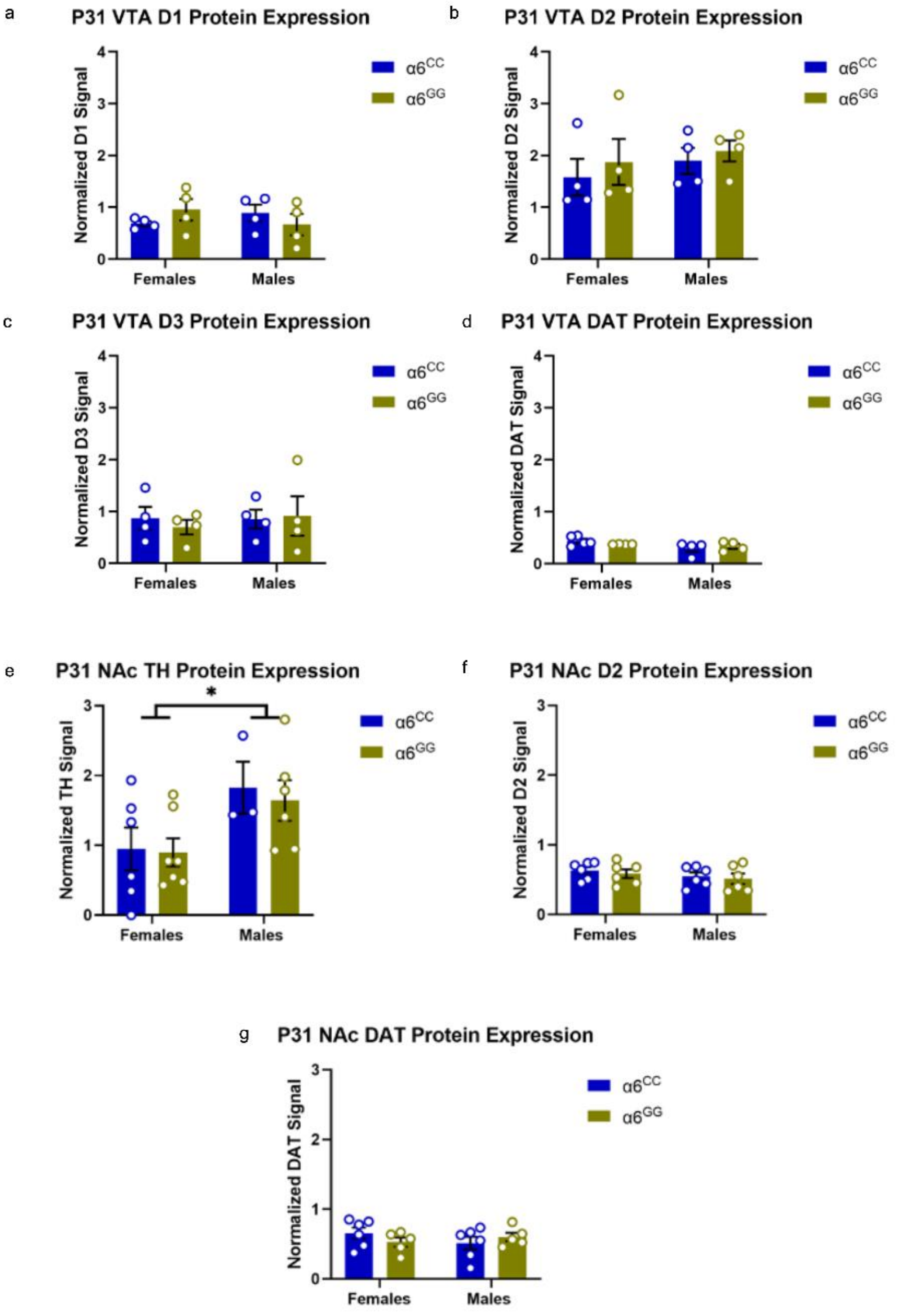
a
P31 VTA D1 Protein Expression
Normalized D1 Signal
Females
Males
α6CC
α6GG
b
P31 VTA D2 Protein Expression
Normalized D2 Signal
c
P31 VTA D3 Protein Expression
Normalized D3 Signal
d
P31 VTA DAT Protein Expression
Normalized DAT Signal
e
P31 NAc TH Protein Expression
*
Normalized TH Signal
f
P31 NAc D2 Protein Expression
Normalized D2 Signal
g
P31 NAc DAT Protein Expression
Normalized DAT Signal

**Supplementary Figure S1. The sex-by-genotype effect on tyrosine hydroxylase is specific to TH and to the VTA.**

**(a–d)** Adolescent (P31) *CHRNA6* 3′-UTR SNP rats show no sex or genotype differences in **(a)** D1, **(b)** D2, **(c)** D3, or **(d)**DAT protein expression within the VTA. n = 4–5 per group. **(e–g)** Dopamine-related protein expression in the nucleus accumbens. **(e)** A two-way ANOVA identified a main effect of sex for TH protein expression ($F(1,17) = 6.06$, $*P = 0.025$), with adolescent males showing higher expression than females independent of genotype. No sex or genotype differences were detected in **(f)** D2 or **(g)** DAT protein levels. n = 3–7 per group. Protein expression was measured by western blot and normalized to GAPDH. Blue denotes $\alpha6^{CC}$ and olive $\alpha6^{GG}$. Bars show mean ± SEM with individual animals overlaid; data points represent independent biological samples. $*P < 0.05$.
